# Structure-Unbinding Kinetics Relationship of p38α MAPK Inhibitors

**DOI:** 10.1101/510255

**Authors:** Xiaoxia Ge, Hepan Tan, Lei Xie

**Affiliations:** School of Medicine, Nankai University, Tianjin, 300071, China; Jixun Computational Solutions, LLC., White Plains, NY, 10601, USA; Department of Computer Science, Hunter College, The City University of New York, New York, NY 10065, USA; The Graduate Center, The City University of New York, New York, NY 10016, USA

**Keywords:** drug efficacy, drug residence time, unbinding kinetics, unbinding pathway, MD simulation, metadynamics, p38α MAP kinase inhibitor, molecular recognition

## Abstract

Rational Drug Design still faces a major hurdle for the prediction of drug efficacy *in vivo* solely based on its binding affinity for the target *in vitro*. The traditional perspective has proven to be inadequate as it lacks the consideration of essential aspects such as pharmacokinetics and binding kinetics in determining drug efficacy and toxicity. Residence time, the average lifetime of drug-target complex, has gained broader recognition as a better predictor for lead optimization. Long residence time could contribute to sustained pharmacological effect and may mitigate off-target toxicity as well. To unravel the underlining mechanism for variation of residence time and determine the ligand features governing the unbinding kinetics, unbinding kinetics of two distinct type II inhibitors of p38α MAP kinase were investigated and compared by molecular dynamics and metadynamics simulation approaches. Free energy landscape of key motions associated with unbinding was constructed for both inhibitors. Multiple unbinding pathways and rebinding were revealed during the drug-target dissociation process of faster unbinder Lig3 and slower unbinder Lig8 respectively, suggesting a novel mechanism of unbinding kinetics. This comparative study implies that hydrophobic and hydrogen-bonding interactions in the R1 group of ligands are crucial for slow unbinding. Such kind of structure-kinetics relationship approaches could also be applied to predict unbinding pathways and kinetics of many other small molecules, and facilitate the design of efficient kinase inhibitors.

## 1. Introduction

Molecular recognition is central to most biological processes. Among many types of molecular recognitions, drug–target recognition has been studied extensively due to their importance in understanding the mechanism of drug action and consequently improving the drug development process.^1^ Equilibrium binding metrics obtained from *in vitro* assays include the half-maximal inhibitory concentration (IC50), the equilibrium dissociation constant (*K*_d_) and the inhibition constant (*K*_i_), *etc*. However, *in vivo* pharmacology often deals with open, non-equilibrium conditions, in which cases equilibrium assumption of drug–target interactions become invalid. Therefore, lead optimization solely based on improving the drug-target binding affinity does not necessarily improve drug efficacy *in vivo*. Residence time, defined as the lifetime of the drug-target complex, is a more relevant *in vivo* metric.^2^ Because long residence time leads to sustained pharmacology, drugs can deliver desirable efficacy at low doses. Lower doses may mitigate off-target toxicity and therefore reduce adverse drug effect (side effect). A static view of the drug-target interaction is inadequate to explain the impact of conformational dynamics on the binding and unbinding processes. For high-affinity interactions, both binding and unbinding processes involve conformational changes of the target.^3^ Conformational change associated with ligand binding/unbinding pathway is an important aspect in understanding the kinetic process.

In the past decade, a number of examples were reported on successful design of new drugs aiming to improve the residence time such as Triazole-Based InhA Inhibitors,^4^ CDK8 inhibitors,^5^ and series of reversible covalent inhibitors.^6–7^ The prediction of kinetic rates based on residue coarse-grained normal mode analysis has been developed and tested successfully for HIV protease inhibitors.^8^

Molecular dynamics (MD) simulation provides a view of the atomic motion over time. It has been a powerful tool for studying biological systems and provided many insights in different areas of research.^9–10^ Drug residence time spans from nanoseconds to hours in time scale. MD simulation results are meaningful only if the trajectories are long enough for the system to visit all the energetically relevant configurations. Practically, some biologically important configurations are often separated by high free-energy barriers; or the system diffuses extremely slowly rendering simulations to be beyond the limit of current computer resources. To tackle these issues, enhanced sampling techniques have been developed in the past few decades including replica exchange, ^11^ accelerated MD,^12^ umbrella sampling,^13^ steered MD,^14^ and metadynamics.^15–17^ These methods facilitate the crossing of free energy barrier via biasing system potential energy or biasing a few selected degrees of freedom, *i.e*., collective variables (CVs). Multiple review articles appeared recently on computational approaches for drug-binding/unbinding kinetics.^18–19^ Metadynamics reconstructs the free energy surface (FES) as a function of the selected CVs using a history-dependent bias potential. Because of its broad applications in thermodynamic and kinetic problems,^20–21^ metadynamics approach has gained increasing attentions.

Protein kinases participate in a number of cellular signaling pathways by catalyzing the phosphorylation of specific substrates, whereby serving as an important family of drug targets. The catalytic activity of a kinase is regulated by its conformation plasticity, *i.e*., it switches between active and inactive forms.^22^ P38 mitogen-activated protein kinases (MAPK) are a class of signal transduction mediators that play pivotal roles in inflammation, cell cycle, apoptosis, cell growth, cell differentiation, senescence and tumorigenesis depending on cell types.^23^ Among four MAPK isoforms p38α, β, γ, and δ, the major one p38α has been identified as drug targets for various inflammatory diseases, cancers, and cardiovascular diseases.^24–26^ Consequently, p38α MAPK has been one of the mostly studied kinases with high-resolution crystal structures available for both the native and type II inhibitor-bound states.^27^ The crystal structure consists a common ATP-binding pocket and a novel allosteric binding site for a class of highly potent and selective diaryl urea inhibitors against human p38α MAPK (Figure 1). The formation of this binding site requires a large conformational change of conserved Asp168-Phe169-Gly170 (DFG) motif not observed previously for any of Ser/Thr kinases. Solution studies demonstrate that this class of compounds has slow binding kinetics, consistent with the requirement for conformational change. One of the most potent compounds in this series, 1-(5-tert-butyl-2-p-tolyl-2H-pyrazol-3-yl)-3-[4-(2-morpholin-4-yl-ethoxy) naphthalen-1-yl]-urea (BIRB796), has picomolar affinity with human p38α MAPK and low nanomolar inhibitory activity in cell culture.^27–28^

**Figure 1.**
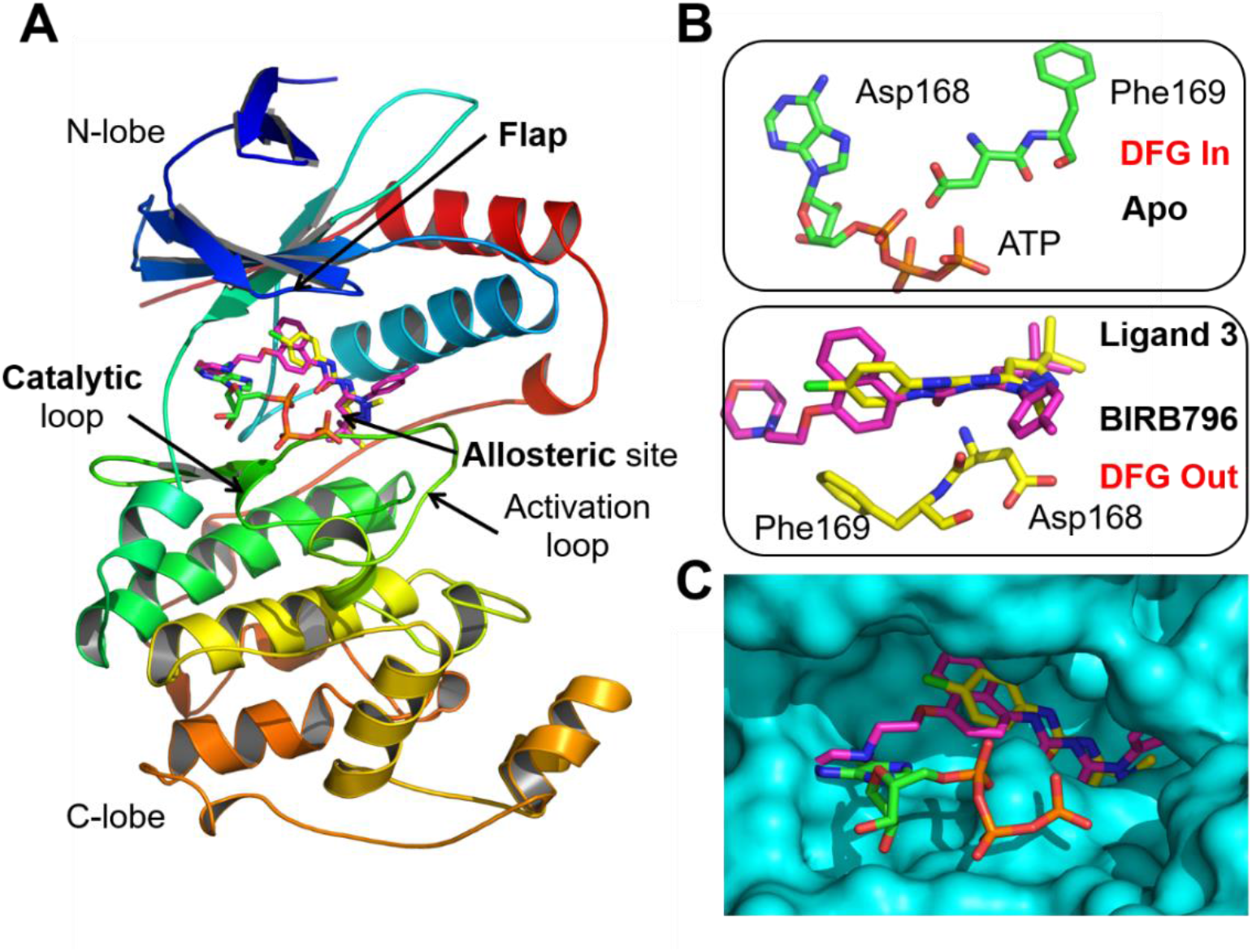
Binding site of the type II inhibitors on p38α MAP kinase. **A)** A schematic view of the structure of p38α in complex with BIRB796 and Lig3. **B)** DFG conformation change upon ligand binding. **C)** Superposition of p38α ligand-binding pocket. The molecular representations were generated with PyMOL^29^ based on PDB structures 1P38, ^30^ 1K1V and 1KV2. ^27^

Quite a few computational studies were carried out in the past few years on this important system, including binding free energy calculations of imatinib, sorafenib, BI-1 and BIRB-796 to p38α.^31^ These studies reveal that the driving force of binding was van der Waals interaction, typically hydrogen bonding.^31^ Sun et al.^32^ found that type II inhibitors unbind through two pathways in opposite directions, *i.e*., allosteric channel and the ATP binding catalytic channel. Casasnovas *et. al*. applied metadynamics and Markov state model to compute the kinetic rates of one inhibitor unbinding from p38α with residence time of a few seconds.^33^ A systematic study of a series of inhibitors spanning a wide range of unbinding kinetics and exhibiting diverse structural features could provide further guidance to therapeutic design. However, it is quite challenging and even prohibited to compute the kinetic rates accurately for complex systems and slower unbinding processes with currently available methodologies and compute resources. As a result, few studies have been performed to compare the unbinding processes of different kinase inhibitors, especially in terms of kinetics. A recent computational study suggested unbinding pathways of p38α’s type-I inhibitors SB2 and SK8, type-II inhibitor BIRB796, and type-III inhibitor LIG4 without ranking order by energetics and kinetics.^34^

Here, we report a computational study of two distinct p38α-BIRB796 analogs regarding their unbinding kinetics and pathways (Table 1). In the present study, we used MD simulation to study conformational dynamics and kinetics during dissociation processes of two drug-target complexes. We then performed metadynamics simulations to improve sampling of slow motions associated with ligand unbinding from p38α. The binding rates of two BIRB796 analogs are within the same order of magnitude, which simplifies the unbinding kinetic problem to a thermodynamic one. Our computational study provides a correct ranking order of the unbinding kinetics that is in agreement with the experimental results, *i.e*., slower unbinding of Lig8 relative to Lig3.^27–28^ Metadynamics simulation of p38α-ligand unbinding is most efficient using two CVs: ligand-cavity distance as CV1 and another geometric parameter as the second CV. The unbinding processes of the two ligands exhibit different FE landscapes of key motions. Slow motions during unbinding include relative orientation between the ligand and the protein, and a distance between the DFG motif and the cavity. Free energy landscape with DFG-cavity distance as CV indicates an inward motion of DFG motif occurring after dissociation of both ligands although they have different FE minima when bound. Rebinding of Lig8 to the secondary binding site during the dissociation process was revealed from the simulations, which is a plausible cause of prolonged residence time for Lig8. The fact of multiple unbinding pathways accessible for Lig3 indicates a higher probability and hence a faster rate of unbinding. Hydrophobic and hydrogen-bonding interactions between the ligand and the protein become unstable and are further disrupted upon unbinding while the binding site undergoes re-solvation. After the ligand unbinds from p38α, the hydrogen bond (H-Bond) network resets in the binding site, as shown in the unbound p38α structure. The embodied methods in this work can be considered as a general approach to understand unbinding kinetics and pathways of other kinase inhibitors of different kinase families. The novel mechanism of unbinding kinetics and multiple pathways revealed from this study can also potentially guide the kinase drug discovery.

**Table 1.**
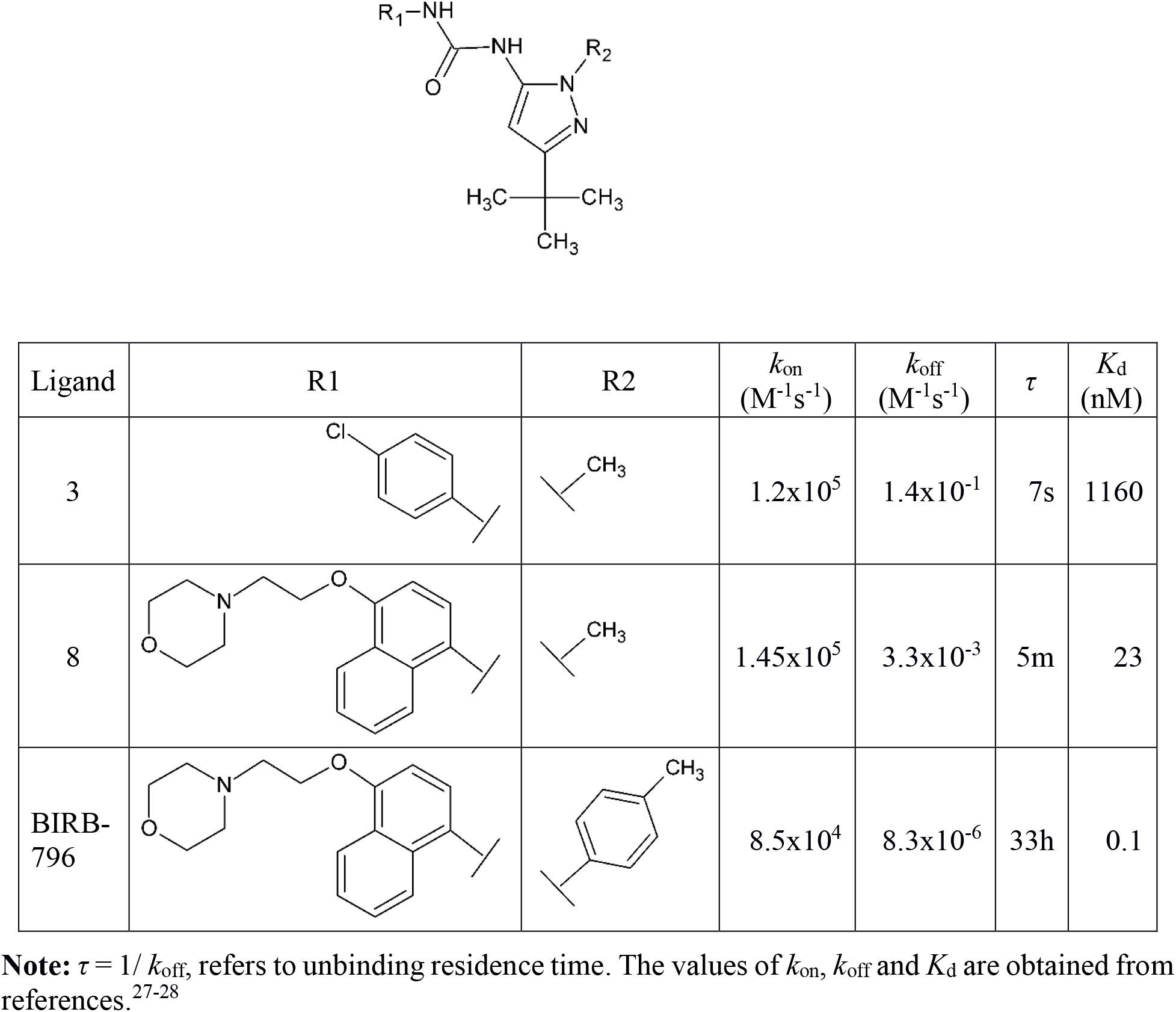
Kinetic and Thermodynamic Unbinding Parameters of p38α inhibitors

| Ligand | R1 | R2 | $k_{\text{on}}$<br>($\text{M}^{-1}\text{s}^{-1}$) | $k_{\text{off}}$<br>($\text{M}^{-1}\text{s}^{-1}$) | $\tau$ | $K_{\text{d}}$<br>(nM) |
| --- | --- | --- | --- | --- | --- | --- |
| 3 | | | $1.2 \times 10^5$ | $1.4 \times 10^{-1}$ | 7s | 1160 |
| 8 | | | $1.45 \times 10^5$ | $3.3 \times 10^{-3}$ | 5m | 23 |
| BIRB-796 | | | $8.5 \times 10^4$ | $8.3 \times 10^{-6}$ | 33h | 0.1 |

**Note:** $\tau = 1/k_{\text{off}}$ , refers to unbinding residence time. The values of $k_{\text{on}}$ , $k_{\text{off}}$ and $K_{\text{d}}$ are obtained from references.<sup>27-28</sup>

## 2. Results and Discussion

### 2.1 Selections of simulation models and slow motions

Two lead compounds of p38α, Lig3 and Lig8 were chosen for this study considering their distinct structures and kinetic properties available from the experiments. As analogs of BIRB796 inhibitor, Lig3 and Lig8 exhibit the same magnitude of binding rates, therefore, the unbinding kinetic problem is simplified to be a thermodynamic one (Table 1). Like many other enhanced sampling methods such as steered MD,^35^ temperature accelerated MD (TAMD),^36^ metadynamics involves sampling over suitable collective variables (CVs).^37^ Appropriate selection of CVs is not a trivial step, because overcoming hidden barriers for slow motions can be a significant challenge for CV-based free-energy calculations. Previous studies on p38α MAPK provide us a wealth of biological information to compare the choices of CVs with a series of calculations.^24–28^

As a metadynamics simulation progresses, fluctuations of CVs become gradually enhanced to facilitate the escape from stable configurations of the system. Multiple sets of CV combinations were tested and led to different profiles of unbinding kinetics as listed in Table 2. Our study suggests that using only the distance between the ligand and the binding cavity on p38α fails to describe the unbinding process properly, which has also been reported by others recently.^33^ Remarkably, for well-converged metadynamics FE simulations (Figure S1), the choice of two different sets of CVs lead to similar results in terms of thermodynamics and kinetics, which could be considered as a validation of our approach.

**Table 2.**
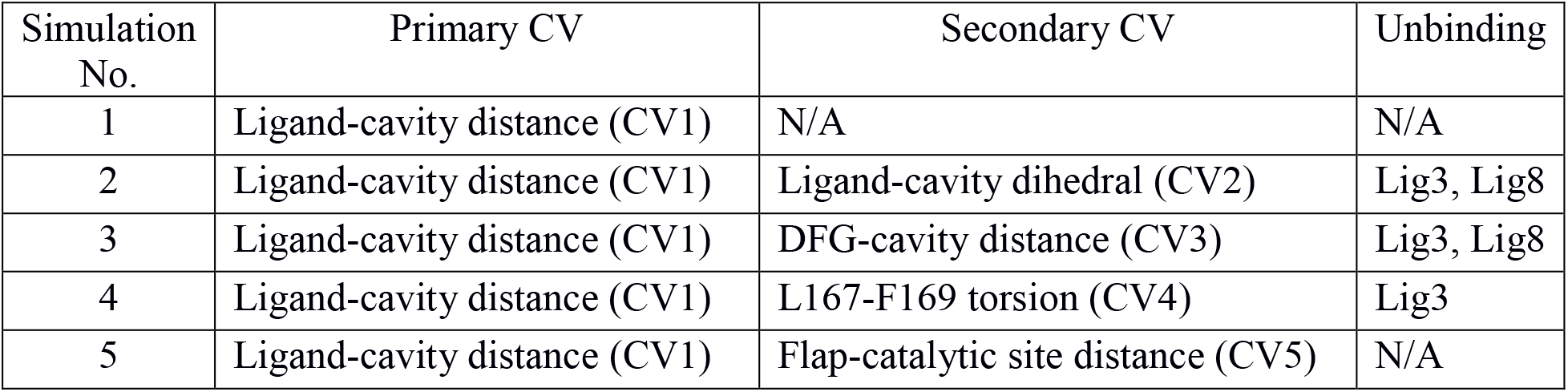
Collective variables (CV) used in metadynamics simulations

### 2.2 Free energy landscape of inhibitor unbinding

Metadynamics simulations using all CV choices CV1-CV2, CV1-CV3, and CV1-CV4 led to consistent results, *i.e*., slower unbinding of Lig8 relative to Lig3 (Figure 2~4). The residence time of ligand unbinding can be defined as τ=1/*k*_off_, where koff is the unbinding rate; the binding constant is defined as *K_b_*= *k*_on_/*k*_off_, where *k*_on_ is the binding rate. Therefore, if inhibitors show similar binding rates, their residence time τ will be proportional to *K_b_*, alternatively reciprocal to the dissociation constant *K_d_*, τ^∝^1/*K_d_*. In another words, stronger binder will bind for a longer time given the same order of magnitude of *k*_on_. The kinetic parameters have been reported for p38α inhibitor BIRB796 and its analogs.^27–28^ In Table 1, inhibitor BIRB796 and its analogs Lig3 and Lig8 all have experimental *k*_on_ of about 1x10^5^ M^−1^s^−1^, however, the three analogous molecules display quite diverse range of residence time from a few seconds to more than a day. In this case, the kinetic variation is solely determined by the binding affinity. Therefore, this unbinding kinetic problem can be reduced to a thermodynamic one.

**Figure 2.**
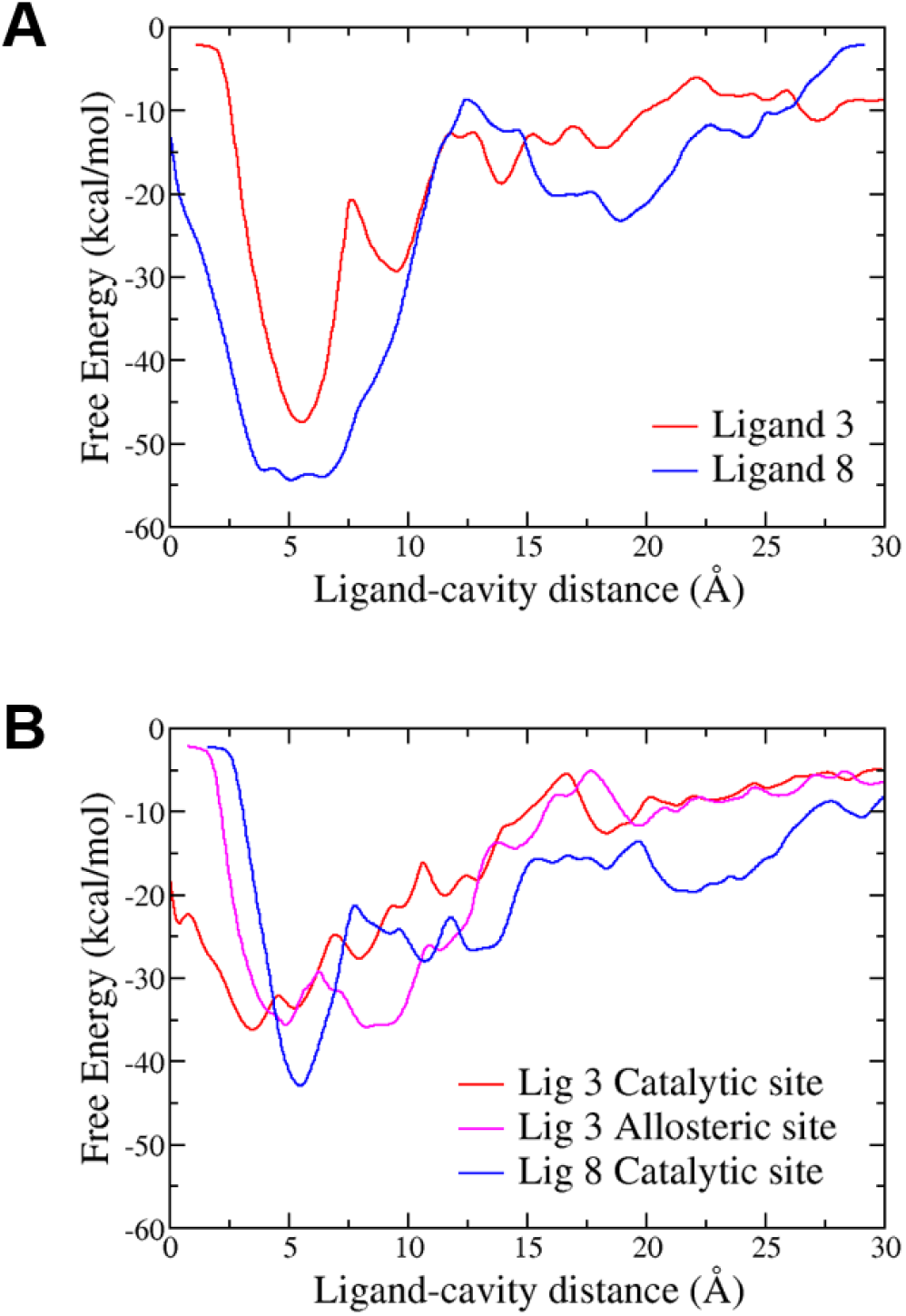
Free energy profiles of p38α inhibitor unbinding. Unbinding FE profiles as a function of CV1 calculated from metadynamics simulation No. 2 (A) and No. 3 (B) respectively. For (B), Ligand 3 adopted different unbinding pathways, *i.e*., through the catalytic site (red) and allosteric site (margenta), while ligand 8 unbinds through the catalytic site (blue) pathway.

**Figure 3.**
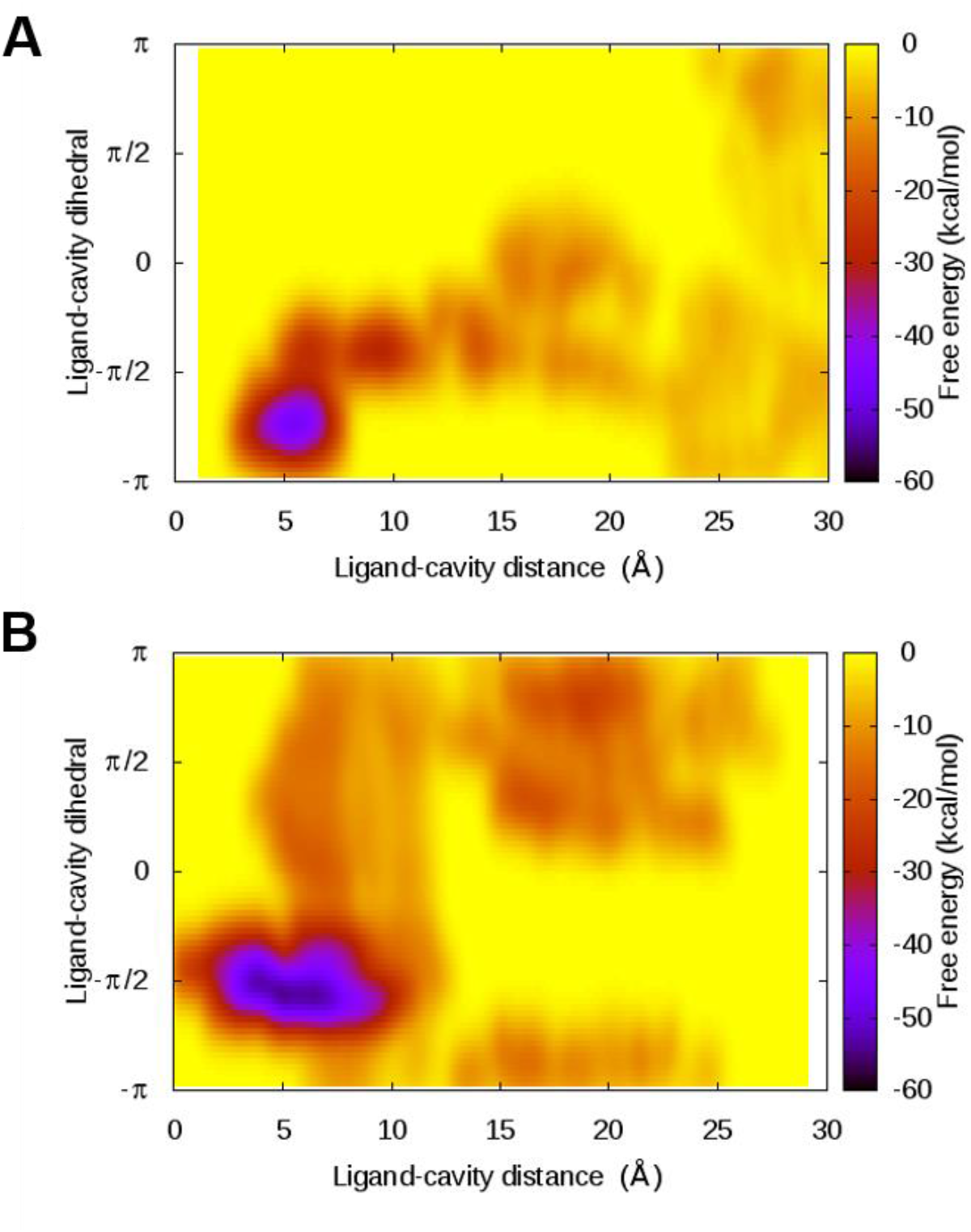
Free energy surfaces of p38α inhibitor unbinding calculated from metadynamics simulations (No. 2) using CV1 and CV2 for ligand 3 (A) and ligand 8 (B).

**Figure 4.**
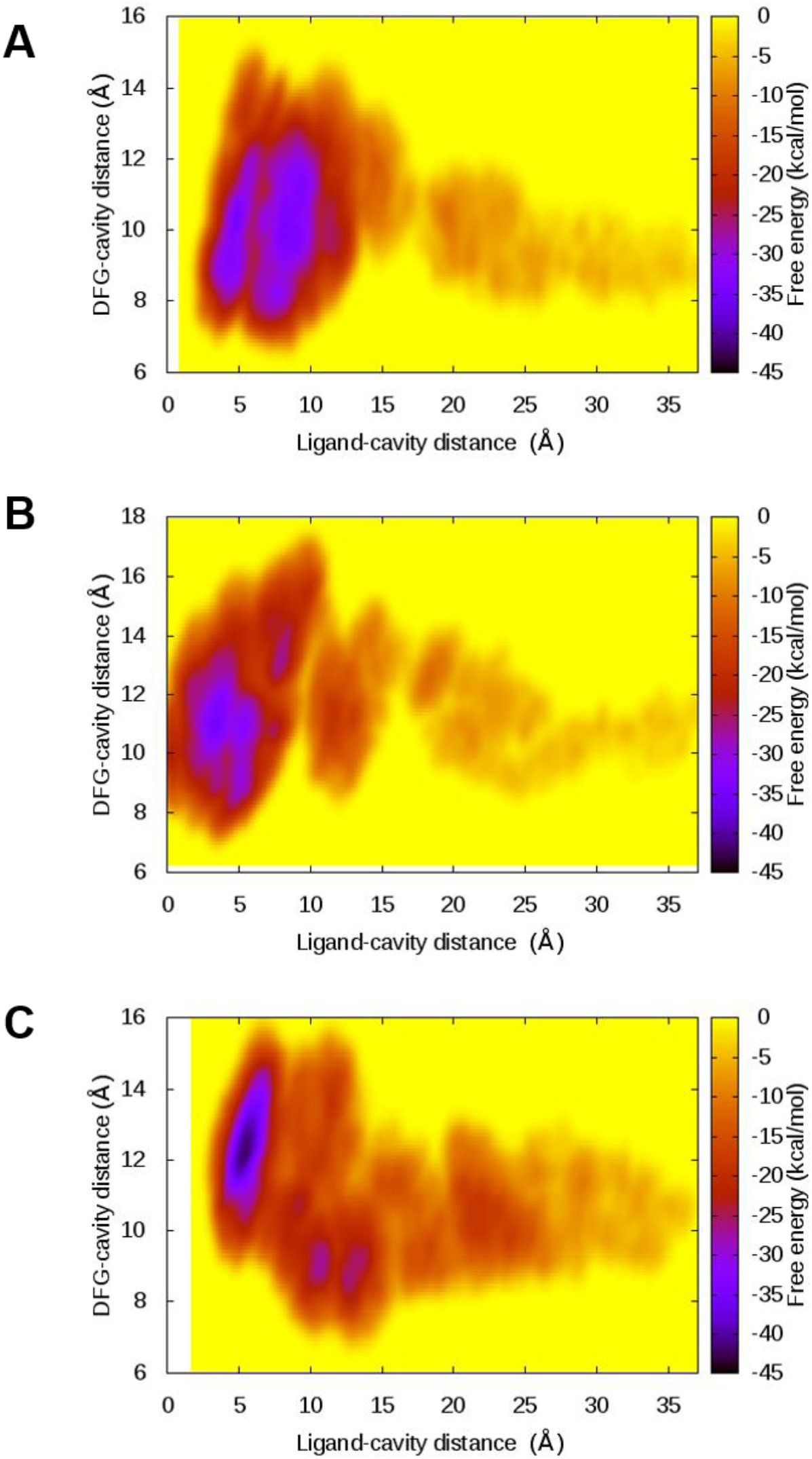
Free energy surfaces of p38α inhibitor unbinding calculated from metadynamics simulations (No. 3) using CV1 and CV3. Ligand 3 unbinds through the allosteric site (A) or the catalytic site (B) and, ligand 8 unbinds through the catalytic site (C).

Unbinding FE profile as a function of ligand-cavity distance calculated based on metadynamics simulations using CV1-CV2 (Figure 2A) suggests that Lig8 has a lower and broader global FE minimum compared to Lig3. Metadynamics simulations using CV1-CV3 (Figure 2B) also confirm that Lig8 has a lower global minimum at the binding cavity in comparison with Lig3. Free energy profiles confirm that Lig8 is a stronger binder of p38α than Lig3. Given similar binding rates,^38^ FE results also indicate that Lig8 unbinds from p38α more slowly than Lig3 does, which is consistent with the experiments.^27–28^

The converged FES reconstructed from metadynamics simulation reveals more information in terms of energetics corresponding to the fluctuation of CVs during the course of unbinding (Figure 3 and 4). To consider the relative orientation between the ligand and the protein, CV2 was defined according to the concept of dihedral angle in previous MD simulation study on ligand-receptor binding.^10, 39^ Using ligand-cavity dihedral as a secondary CV, unbinding free energy landscape of Lig8 is more complex than that of Lig3. The FE map demonstrates that Lig3 has a narrower FE minimum of about −47 kcal/mol in the binding cavity when CV1 is about 6 Å and CV2 is about −140 ° (Figure 2A and 3A). By contrast, Lig8 has a broader binding FE minimum of about −54 kcal/mol for Lig8 when the dihedral is about −90° with ligand-cavity distance spanning from 3 Å to 8 Å (Figure 2A and 3A). Upon unbinding, Lig8 start to change its conformation and orientation relative to the protein while remain in the binding cavity with a CV1 of ~5 Å. Using DFG-cavity distance as the second CV, the FES exhibits similar characteristics in metadynamics simulations with CV1-CV3, as in simulations with CV1-CV2 (Figure 4). The binding of type II inhibitors Lig3 and Lig8 both lead to outward orientation of DFG motif. The DFG-cavity distance (CV3) fluctuates between 8 Å and 12 Å when Lig3 bound and increases to 11~14 Å when Lig8 is in the cavity.

### 2.3 Unbinding kinetics and rebinding

In the equilibrated configuration, the distance between the inhibitor and the cavity fluctuates between 5 Å and 10 Å (Figure 5~7), the inhibitor is considered fully unbound when CV1 is greater than 20 Å. For simulation using CV1-CV2, Lig3 unbinding starts at ~1 ns (Figure 5A and 5C), while for Lig8, unbinding occurs after 2.5 ns followed with rebinding at 3.5 ns before complete dissociation (Figure 5B and 5D).

**Figure 5.**
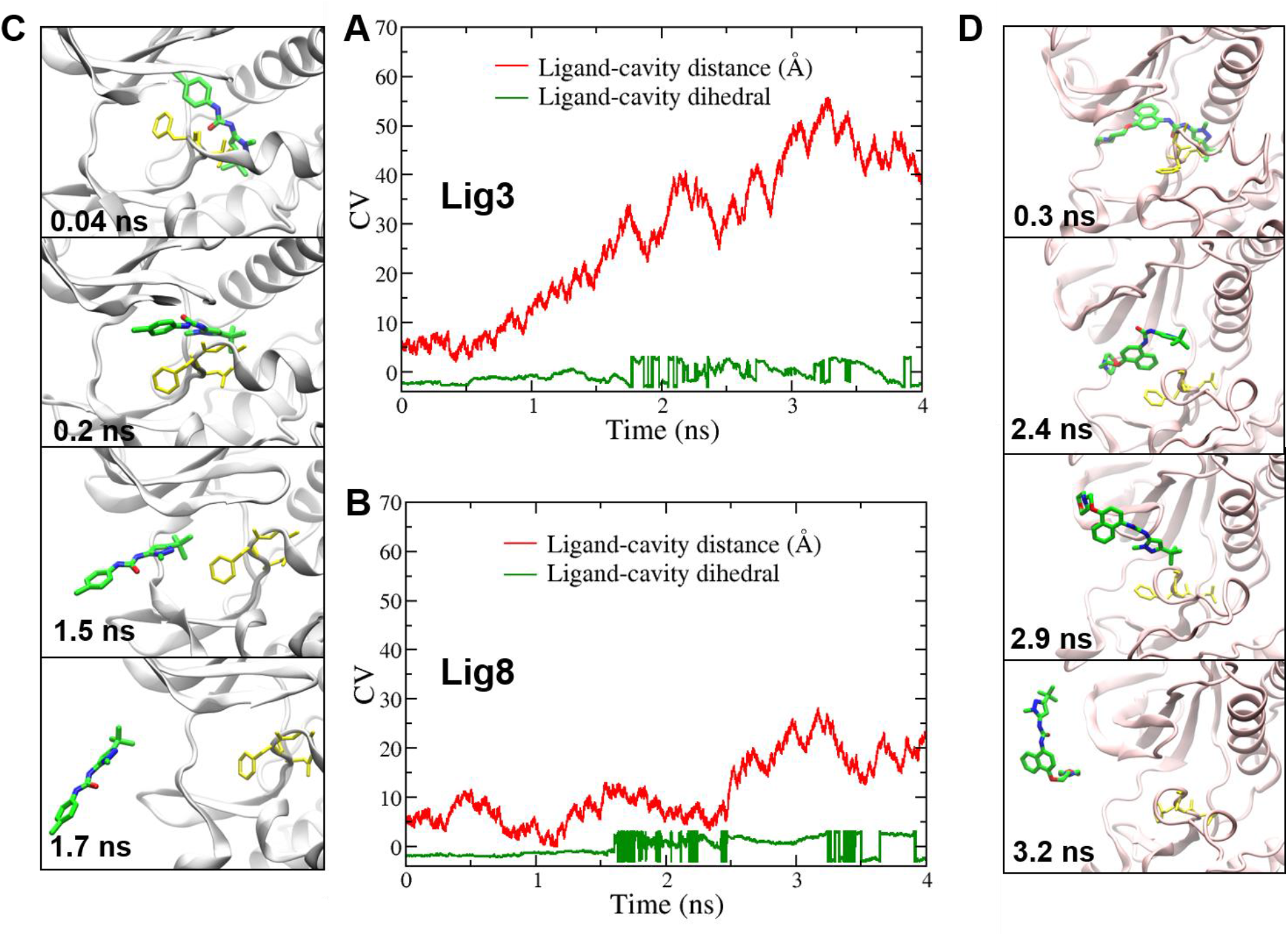
Metadynamics simulations using CV1 and CV2. Time series of collective variables CV1 (red) and CV2 (green) are shown for ligand 3 (A) and ligand 8 (B) with representative snapshots taken from the trajectories for ligand 3 (C) and ligand 8 (D) respectively. In (C) and (D), snapshot conformations are shown for p38α (cartoon), the ligand (colored licorice) and non-hydrogen atoms of DFG loop (yellow licorice). The images of snapshot conformations were created with VMD^40^

**Figure 6.**
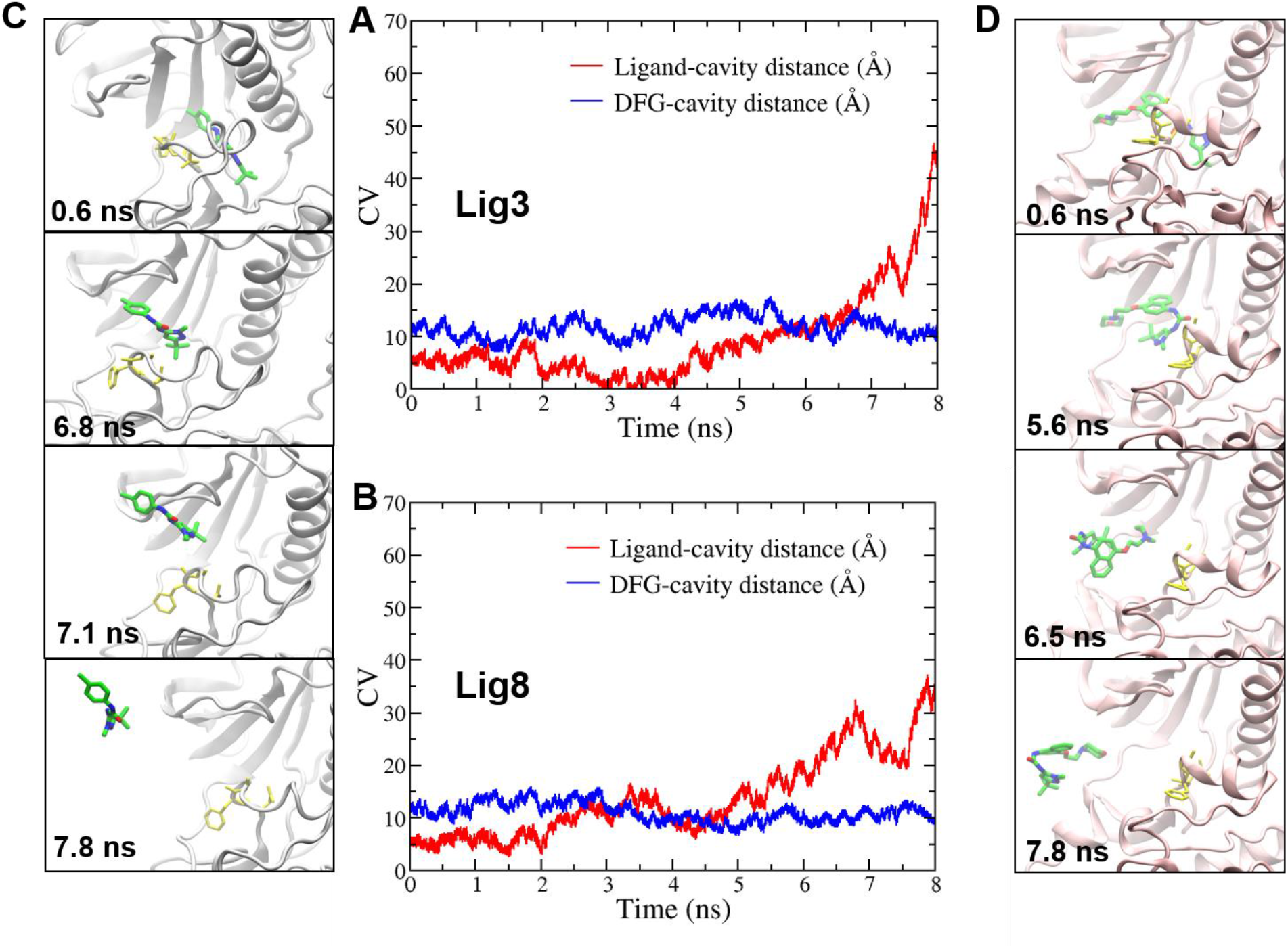
Metadynamics simulations using CV1 and CV3. Time series of collective variables CV1 (red) and CV3 (blue) are shown for ligand 3 (A) and ligand 8 (B) with representative snapshots taken from the trajectories for ligand 3 (C) and ligand 8 (D) respectively.

**Figure 7.**
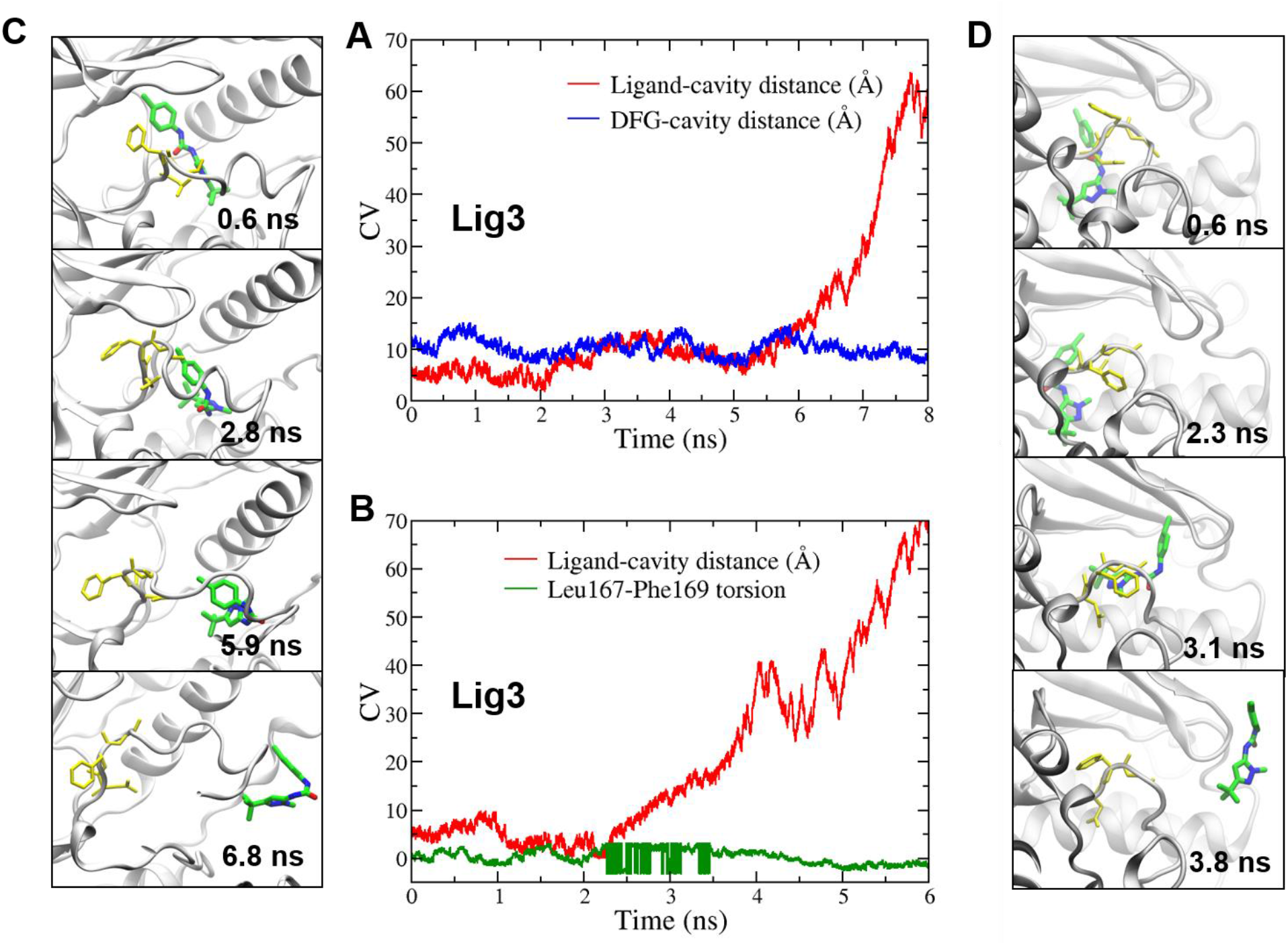
The alternative unbinding pathways for ligand 3. Time series of collective variables for metadynamics simulations using CV1-CV3 (A) and CV1-CV4 (B) with representative snapshots taken from trajectories using CV1-CV3 (C) and CV1-CV4 (D) respectively. (A) and (C) represent unbinding through the allosteric site, while (B) and (D) suggest the third unbinding pathway from the P-loop. The CVs in time series are colored in red, blue and green for CV1, CV3 and CV4 respectively.

In metadynamics simulations using CV1-CV3, Lig3 unbinding occurs after 6 ns (Figure 6A and 7A), while for Lig8, this occurs at about 6 ns and the ligand rebinds at 7.5 ns before complete dissociation from the protein (Figure 6B and 6D). In the meantime, the protein exhibits different FE landscapes of key motions during unbinding. Rebinding to the secondary binding site underneath the glycine-rich phosphate-binding loop (P-loop) also potentially contributes to the longer residence time of Lig8 (Figure 5B and 6B). The R1 group ethoxy-morpholine-naphthalene on Lig8 is bulkier than the chlorophenyl ring on Lig3 (Table 1). The naphthalene ring is more lipophilic than phenyl ring, while morpholine could form H-Bond with nearby Met109.^38^ All these interactions strengthen the binding affinity and decrease the unbinding rate of Lig8. Lig8 rebinds to p38α surface through interaction between this functional moiety and Val30 and Val38 side chain on the P-loop (Figure 5D and 6D). The rebinding of Lig8 leads to prolonged apparent target occupancy, and consequently longer residence time.

### 2.4 Multiple unbinding pathways

In this study, metadynamics simulations of Lig8 unbinding using different sets of CVs result in a single pathway through the ATP-binding site, also known as the catalytic channel (Figure 5D and 6D).^32^ By contrast, Lig3 unbinding exhibits three different unbinding pathways: 1) through the catalytic site (Figure 5C and 6C), 2) through the allosteric site (Figure 7C), and 3) opening the glycine-rich P-loop (Figure 7D). The common feature of Lig3 unbinding through the catalytic pathway and the flap pathway is that inhibitors exit the binding site with their R1 aromatic-ring group exposed to the solvent first (Figure 5D, 6D and 7D). In comparison, Lig3 unbinding through the allosteric pathway requires complex conformational changes (Figure 7C). During the simulation, Lig3 became unstable at 2.8 ns, and then oriented itself into a bended conformation and began to egress the binding cavity at 5.9 ns, finally fully escaped the receptor at 6.8 ns (Figure 7C).

The correlation of DFG motion with unbinding suggests that CV3, the distance from DFG motif to cavity, is a slow motion relevant to the unbinding through the allosteric pathway. There exists a higher energy barrier for Lig8 to escape the allosteric site. Multiple unbinding pathways available for Lig3 lead to a higher probability and hence a faster rate of unbinding, which suggests a novel mechanism of unbinding kinetics for p38α inhibitors.

### 2.5 The DFG motif motion

The DFG loop regulates kinase activation by altering side chain conformation.^22^ The kinase become inactive by adopting an DFG-out conformation when type II inhibitors are bound.^27^ It was also reported that the backbone of DFG motif functions as a hydrophobic core to interact with the t-butyl tail of the inhibitors.^33^ From metadynamics simulations using CV1-CV3, we found that for both inhibitors, the DFG-cavity distance is shortened by 5~10 Å after the inhibitor unbinding. This ligand-free conformation mimics the active conformation of an apo kinase. Remarkably, before dissociation of ligands, a quick inward motion of ~5Å were also observed for DFG motif relative to the cavity (Figure 6A, 6B and 7A). Then facilitated by the DFG outward motion, the cavity opened to allow the ligand to exit, as indicated by the increase of DFG-cavity distance (10 Å to ~15 Å). After ligand dissociation, DFG-cavity distance was shortened to ~10 Å. Therefore, FE landscape with the DFG-cavity distance as the second CV suggests that the dynamic motion of DFG motif is coupled with the dissociation of both ligands, even though they have different FE minima when bound.

### 2.6 The Leu167-Phe169 side chain torsion change

It was reported that the side chain opening of Phe169 accompanies with BIRB796 inhibitor binding.^38^ During 6 ns metadynamics simulation with the second CV defined by LF side chain torsion Leu167Cβ-Leu167Cα-Phe169Cα-Phe169Cγ (CV4), only Lig3 unbinds at ~3 ns, Lig8 remains bound throughout 6 ns simulations (Figure 7B and Figure S2). The kinetic difference implies that using the LF side chain torsion is not sufficient to characterize the unbinding process of Lig8. Therefore, CV4 is not an appropriate choice of the second CV. This demonstrates the nature of complex conformation dynamics controlling the unbinding kinetics.

Electrostatic and hydrophobic interactions stabilize ligand-binding to p38α binding site. Residue Lys53, Asp168 and Arg173 form H-Bonds with water in the binding site when Lig3 is bound (Figure S3 top). Two urea amide groups of the ligand form H-Bonds with the side chain carboxylate group of Glu71. Upon Lig3 dissociation, H-Bond interactions between the ligand and the protein become unstable and further disrupted (Figure S3 middle). In the meantime, the binding site undergoes re-solvation. After Lig3 unbinds from p38α, the H-Bond network resets in the binding site (Figure S3 bottom), resembling the ligand-unbound p38α structure.

By comparing the ligand structures in Table 1, all three compounds have a common scaffold of a pyrazole ring attached to a urea group. Lig3 bears a benzenyl chloride at R1 position, while Lig8 bears an ethoxy morpholin attached to naphthalene at R1 position, rendering extra hydrophobicity to bind p38α more strongly in the hydrophobic cavity underneath the P-loop.

### 2.7 The P-loop (Flap) motion

A glycine-rich motif on the P-loop forms a flexible flap at the catalytic site.^41^ To monitor the open-and-close motion of the flap and characterize its relationship with inhibitor unbinding, another collective variable CV5 was defined as a distance between Cα atoms of Ser32, Gly33, Ala34 and Thr35 and Cα atoms of the catalytic site residues Asn155, Leu156, and Ala157. However, neither Lig3 nor Lig8 is able to disassociate from p38α within 8 ns of metadynamics simulations using CV1 and CV5 (Figure S4). Therefore, the flap motion is not a slow motion directly related to ligand unbinding kinetics.

Although current study generates correct rank order of binding affinities and unbinding rates for the two ligands, a fairly large discrepancy of the absolute FE values remains an issue. As a fast-developing CV-biasing method, the limitation of metadynamics-based methods should not be neglected. When using these methods, one should pay more attention on the efficiency and convergence, especially potential mis-interpretation of converged FES calculation over sub-optimized reaction coordinate.^42^ Nevertheless, for such kind of systems with high therapeutic values, future study should be focused on optimizing CVs and applying more efficient and comprehensive algorithm to the sampling.

## 3. Materials and Methods

### 3.4 System setup

The kinetics of BIRB796 analogs binding to p38α MAPK have been measured experimentally as listed in Table 1.^27–28^ High-resolution 3D structures of p38α MAPK are abundant in the Protein Data Bank. Crystal structure of p38α MAPK in complex with Lig3 (PDB code 1KV1) was used for the initial conformation of the protein. All simulation models were constructed as monomers encompassing residue number 5 to 352. Missing residues in the flap and the activation loop were added using the FALC-Loop server.^43^ The loop conformations were generated by fragment assembly and analytical loop closure. The model p38α MAPK in complex with Lig8 was built by modifying the ligand structure in 1KV1.

### 3.2 Equilibration with MD simulation

All simulations were carried out using NAMD 2.8 package^44^ patched with PLUMED 1.3^45^. CHARMM36 all-atom force field^46^ was used for protein and salt ions. CGenFF force field parameters for BIRB796 analogs were created with ParamChem and validated using Gaussian09.^47^ The systems were first solvated with a box of about 17500 TIP3P water molecules,^48^ neutralized with 56 Na+ and 49 C1- ions to reproduce the ionic strength of the experiments.^28^ Periodic Boundary Condition was applied throughout the simulation with a box size of 102 × 80 × 76Å^3^. While keeping protein nonhydrogen atoms fixed, ions and hydrogens were first relaxed by performing 1000 steps of conjugate gradient energy minimization. Then with protein backbone atoms fixed, the systems were relaxed by performing 500 steps of minimization. The system was then subjected to 500 steps of minimization and 20 ps heating with the protein restrained with a 1 kcal/mol/Å^2^ harmonic force constant. After removal of harmonic constraints, the system underwent Langevin dynamics with Nose–Hoover barostat^49–50^ at constant pressure of 1.0132 bar and 298 K temperature. A cutoff of 12 Å and a switching distance of 10 Å were applied for non-bonded interactions. Electrostatics was evaluated with the particle mesh Ewald method. The SETTLE algorithm^51^ was used to constrain all bonds involving hydrogen atoms.

### 3.3 Metadynamics simulations of unbinding

The equilibrated structures were used as starting points for metadynamics simulation. Geometrical collective variables (CV) were defined to characterize the unbinding pathway, as listed in Table 2. Specifically, CV1 is the distance between the centroid of the central diaryl urea group on the ligand and the centroid of Cα atoms of binding cavity residues Leu74, Leu104, and Gly137; CV2 is the dihedral angle between two major inertia axes of the ligand and the protein, identified by two centroids on the ligand (the central diaryl urea group and benzenyl group) and centroids of two sets of Cα atoms in the stable α-helices composed of residue number 131–140 and 204–213).

To study the key motions of the activation loop during ligand dissociation from the protein, additional geometrical CVs were used. CV3 was defined as the distance between the centroid of three Cα atoms of the DFG motif (Asp168, Phe169 and Gly170) and the centroid of three Cα atoms of binding site residues Leu74, Leu104, and Gly137. CV4 were defined to characterize the side chain motion of Phe169 on the DFG motif, *i.e*., torsion angle Leu167Cβ-Leu167Cα-Phe169Cα-Phe169Cγ.

To monitor the open-and-close motion of the flap, a distance CV5 was defined between the centroid of four Cα atoms of Ser32, Gly33, Ala34 and Thr35 and the centroid of three Cα atoms of the catalytic site residues Asn155, Leu156, and Ala157. Using the equilibrated structures, two independent runs of 8 ns simulations were carried out using each of the five CV sets for both ligands (Table 2), with a total of 20 simulations. The multi-dimensional unbinding FE landscapes were reconstructed based on the Gaussians with optimal height and width. In all simulations, the Gaussian-shaped potentials were deposited every 100 simulation steps with an initial height of 0.1 kcal/mol. The Gaussian width was set to 0.35 Å for ligand-cavity distance, 0.25 Å for DFG-cavity distance and flap-catalytic site distance, and 0.35 for ligand-cavity dihedral and Leu167-Phe169 torsion angle. The convergence of the metadynamics simulations was monitored and ensured by calculating the estimate of FES with an interval of every 100 Gaussians deposited.

## 4. Conclusion

In the present study, metadynamics simulations were performed to enhance sampling of slow motions associated with ligand unbinding from p38α MAPK. Two analogs of BIRB796 inhibitor share the same magnitude of binding rates, therefore, the unbinding kinetic problem was simplified to be a thermodynamic one. Current study of type II inhibitors confirms slower unbinding of Lig8 relative to Lig3. Metadynamics simulation of p38α-ligand unbinding is most efficient using two CVs: the ligand-cavity distance as CV1 and another geometric parameter as the second CV. The unbinding kinetics involves different FE landscapes of the key motions. Slow motions related to unbinding are relative orientation between the ligand and the protein, and the distance from the DFG motif to the cavity.

FE profiles confirm that Lig8 binds to p38α more strongly than Lig3. The unbinding pathway of Lig8 is only found to be initiated from the catalytic site; rebinding of Lig8 to the secondary binding site underneath the P-loop may contribute to the longer residence time. The R1 group ethoxy-morpholin-naphthalene group on Lig8 is bulkier than the chlorophenyl ring on Lig3, providing extra hydrophobic and H-Bond interactions with the cavity, and consequently strengthen the binding affinity and decrease the unbinding rate of Lig8. Lig8 rebinds to p38α surface through interaction between this functional moiety and Val30 and Val38 side chain on the P-loop. Other than the previously reported two unbinding pathways, the catalytic channel and the allosteric channel,^34^ an additional unbinding pathway of Lig3 underneath the P-loop were revealed. Multiple unbinding pathways available for Lig3 lead to a higher probability and hence a faster rate of unbinding, which suggests an underlying mechanism of unbinding kinetics for p38α inhibitors.

According to FES with DFG-cavity distance as the second CV, inward motion of DFG motif occurs upon dissociation of both ligands, albeit with different FE minima in ligand-bound state.

Although the P-loop in the p38α catalytic site is important for ligand unbinding, the flap motion itself is not a slow motion that is directly associated with unbinding. Hydrophobic and H-Bond interactions between the ligand and protein become unstable and further disrupted upon unbinding while the binding site undergoes re-solvation. After the ligand unbinds from p38α, the H-Bond network is recovered in the binding site, as shown in the unbound p38α structure.

To summarize, the approach utilized in our study can be applied to understand unbinding kinetic and pathway of other kinase inhibitors even for different kinase families. The kinetic information obtained from this study can potentially guide the kinase inhibitor design.

## Acknowledgments

This work was partly supported by Grant Numbers R01LM011986 from the National Library of Medicine (NLM), R01GM122845 from the National Institute of General Medical Sciences (NIGMS), and R01AD057555 of National Institute of Aging of the National Institute of Health (NIH). The authors acknowledge the compute resources and support provided by CUNY High Performance Computing Center (HPCC). The CUNY HPCC is operated by the College of Staten Island and funded, in part, by grants from the City of New York, State of New York, CUNY Research Foundation, and National Science Foundation Grants CNS-0958379, CNS-0855217 and ACI 1126113.

## Conflicts of Interest

The authors declare no conflict of interest.

