## Supporting Information for "Structure-Unbinding Kinetics Relationship of p38α MAPK Inhibitors"

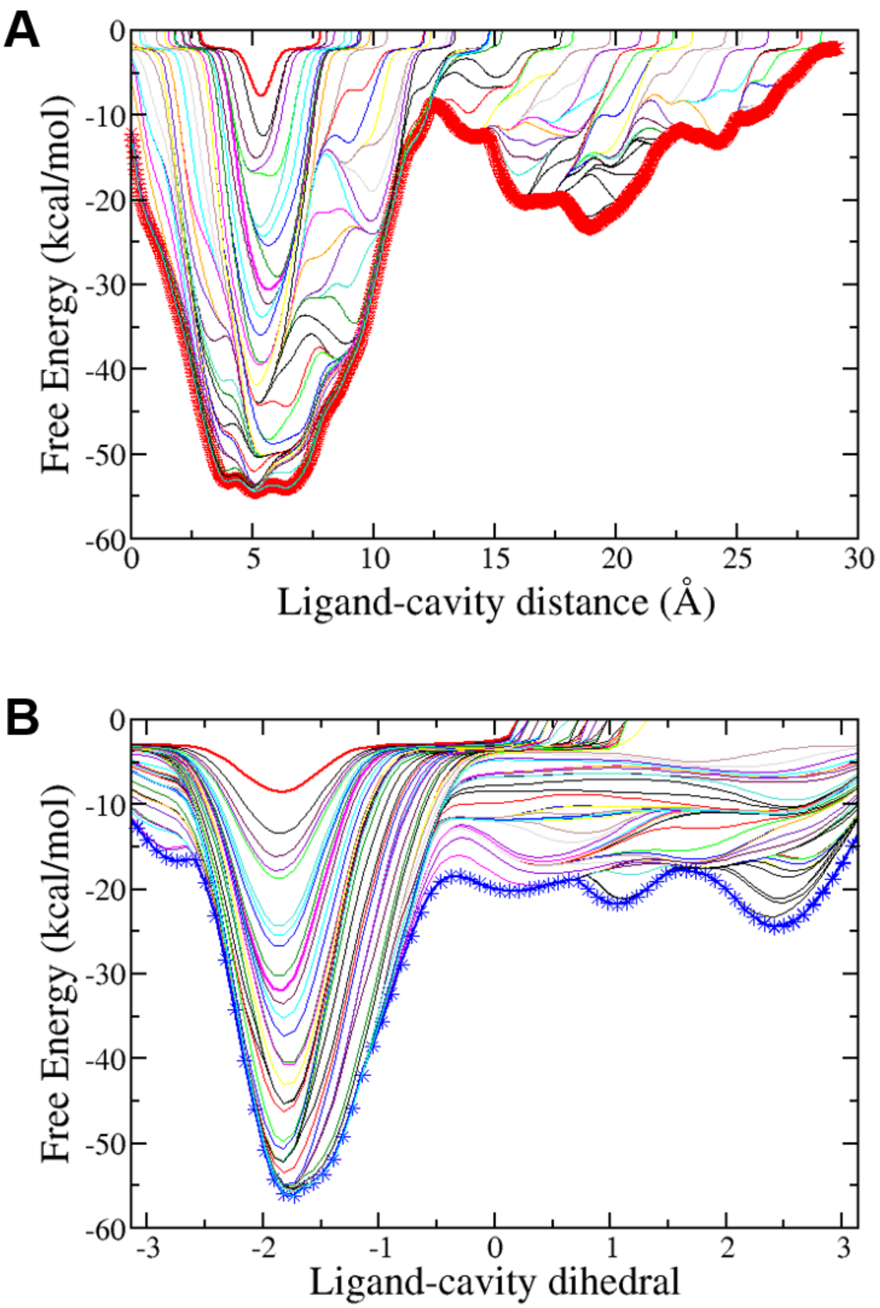


**Figure S1.** Convergence of metadynamics FE calculation using CV1 (A) and CV2 (B) for lig8. Each curve represents a FES profile generated every 100 gaussian hills, the last 500 hills were highlighted by thick curves with crosses.

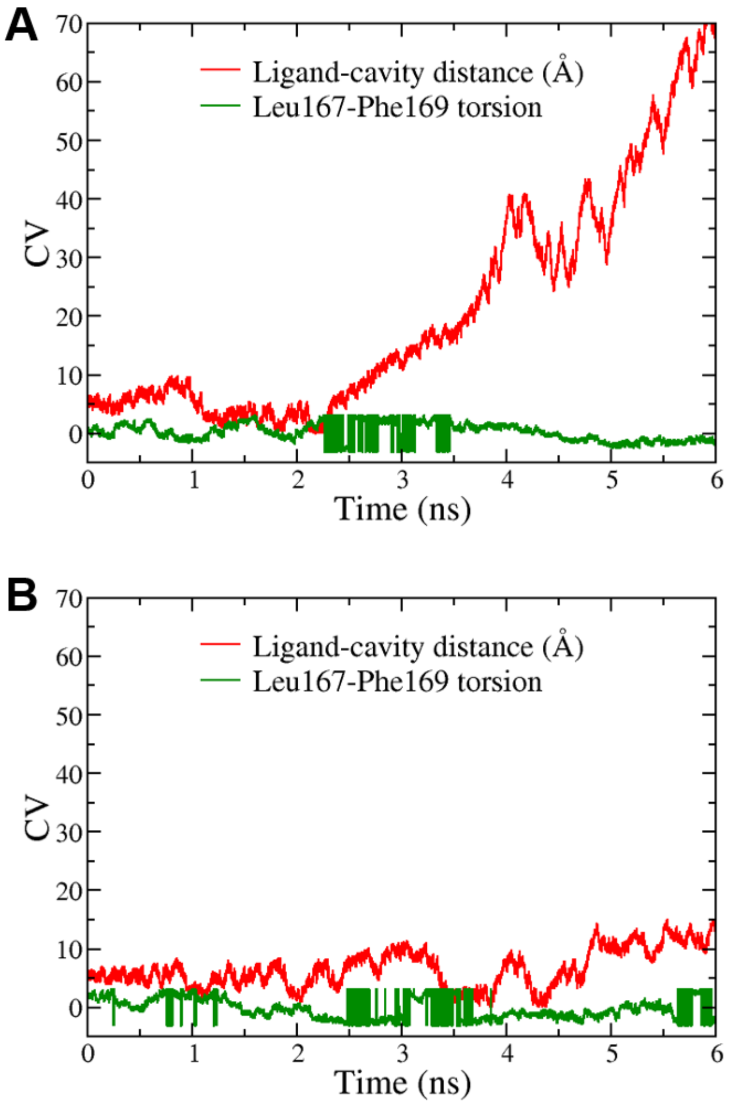


**Figure S2.** Time series of the collective variables in metadynamics No. 4. CV1 and CV2 are colored in red and green respectively for ligand 3 (A) and ligand 8 (B).

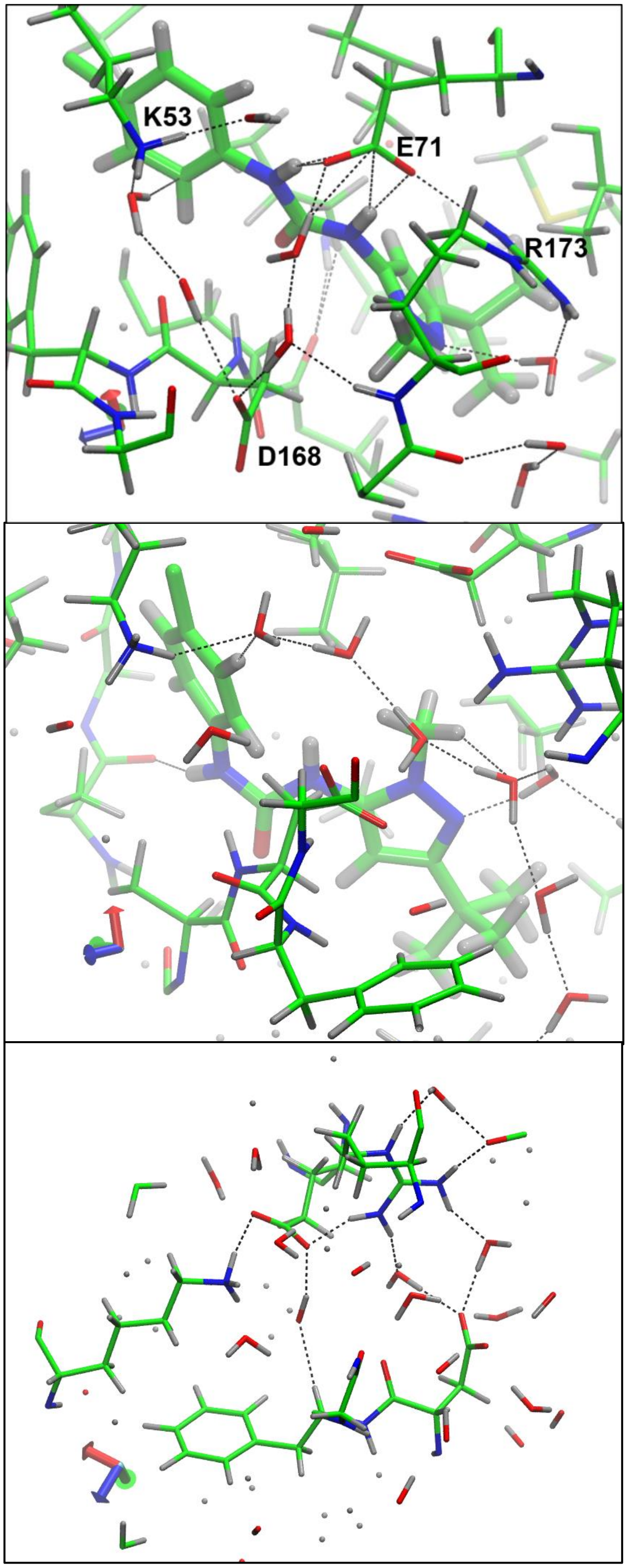


**Figure S3.** Binding site electrostatic interactions during Lig3 unbinding. Snapshots were taken from metadynamics with CV1 and CV4, when ligand is bound (top), starting to unbind at 2.3 ns (middle) and disassociated at 5.9 ns (bottom)

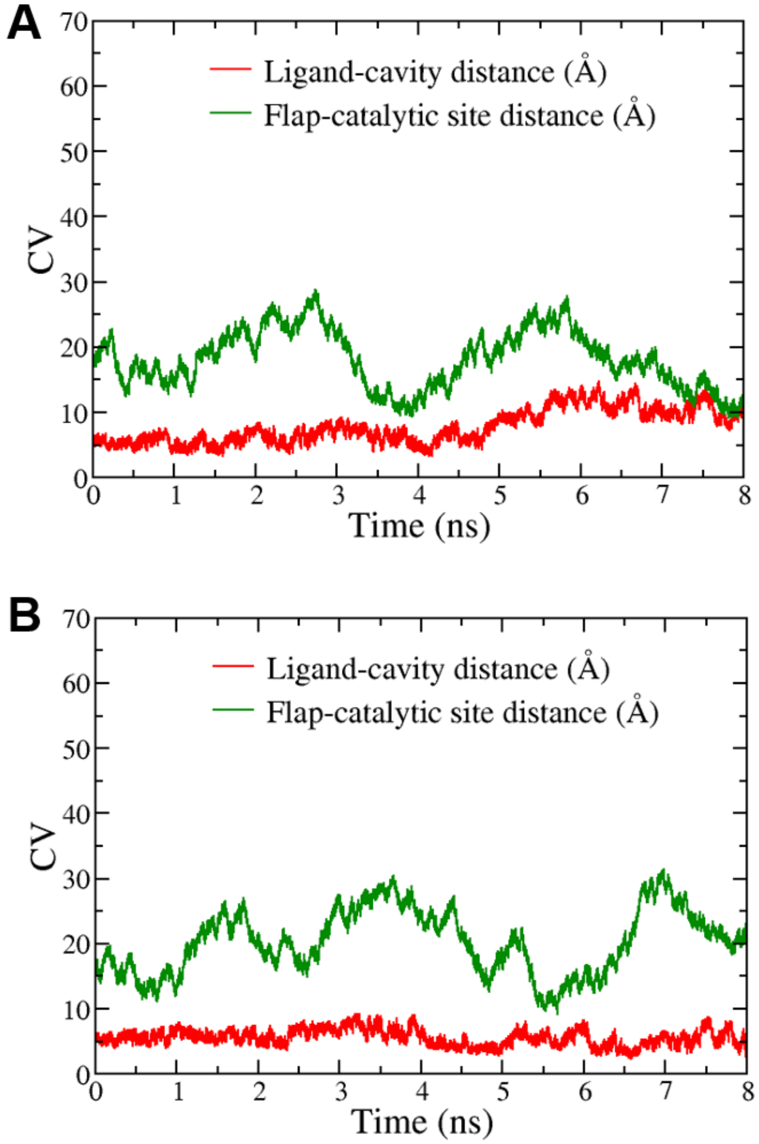


**Figure S4.** Time series of the collective variables CV1 (red) and CV5 (green) for ligand 3 (A) and ligand 8 (B).

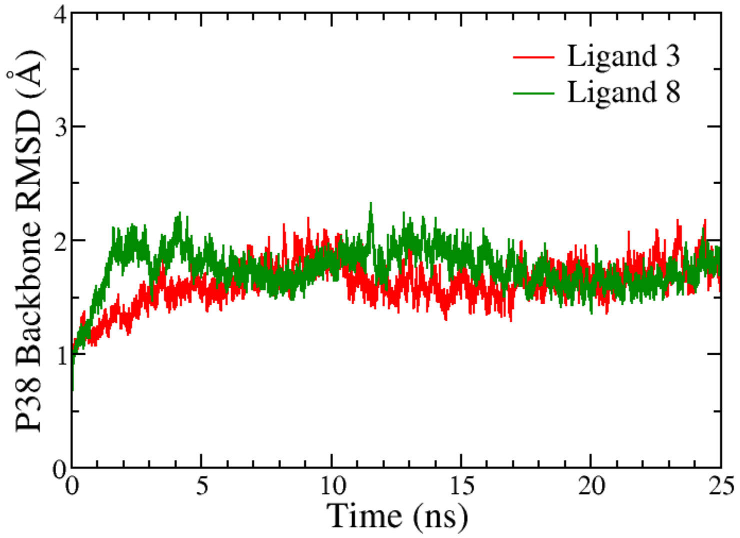


**Figure S5.** RMSD time series of backbone non-hydrogen atoms on p38 kinase in complex with ligand 3 (red) and ligand 8 (green).
